# Limited overlap in RNA virome composition among rabbits and their ectoparasites reveals barriers to virus transmission

**DOI:** 10.1101/812081

**Authors:** Jackie E. Mahar, Mang Shi, Robyn N. Hall, Tanja Strive, Edward C. Holmes

**Author notes:** Corresponding author: Edward C. Holmes, Marie Bashir Institute for Infectious Disease and Biosecurity, Charles Perkins Centre, School of Life and Environmental Sciences and Sydney Medical School, The University of Sydney, Sydney, NSW 2006, Australia.

## Abstract

Ectoparasites play an important role in virus transmission among vertebrates. However, little is known about the extent and composition of viruses that pass between invertebrates and vertebrates. In Australia, flies and fleas support the mechanical transmission of viral biological controls against wild rabbits - rabbit haemorrhagic disease virus (RHDV) and myxoma virus. We compared virome structure and composition in rabbits and these associated ectoparasites, sequencing total RNA from multiple tissues and gut contents of wild rabbits, fleas collected from these rabbits, and flies trapped sympatrically. Meta-transcriptomic analyses identified 50 novel viruses from multiple RNA virus families. Rabbits and their ectoparasites were characterised by markedly different viromes: although viral contigs from six virus families/groups were found in both rabbits and ectoparasites, none were vertebrate-associated. A novel calicivirus and picornavirus detected in rabbit caecal content were vertebrate-specific: the newly detected calicivirus was distinct from known rabbit caliciviruses, while the novel picornavirus clustered with the *Sapeloviruses*. Several *Picobirnaviridae* were also identified, falling in diverse phylogenetic positions suggestive of an association with co-infecting bacteria. The remaining viruses found in rabbits, and all those from ectoparasites, were likely associated with invertebrates, plants and co-infecting endosymbionts. While no full genomes of vertebrate-associated viruses were detected in ectoparasites, suggestive of major barriers to biological transmission with active replication, small numbers of reads from rabbit astrovirus, RHDV and other lagoviruses were present in flies. This supports the role of flies in the mechanical transmission of RHDV and implies that they may assist the spread of astroviruses.

## Introduction

Ectoparasites act as vectors for many notable viral pathogens of vertebrates, including Zika virus, dengue virus, and tick-borne encephalitis virus (Boyer, Calvez, Chouin-Carneiro, Diallo, & Failloux, 2018; Lindquist & Vapalahti, 2008; Rodhain, 2015). Transmission can occur “biologically”, with active virus replication in the ectoparasite, or “mechanically” without ectoparasite replication (Chihota, Rennie, Kitching, & Mellor, 2001; Kuno & Chang, 2005; McColl et al., 2002; Rodhain, 2015). Both mechanisms enable viruses to spread across spatial or ecological barriers that might inhibit direct transmission (Rosenberg & Beard, 2011). Ectoparasites are predominantly arthropods, including such animals as lice and fleas, as well as intermittent ectoparasites such as mosquitos, ticks and blowflies (Hopla, Durden, & Keirans, 1994).

The European rabbit (*Oryctolagus cuniculus*) has been profoundly impacted by ectoparasite-mediated viral transmission. As rabbits are a pest species in Australia, two virus biological controls - rabbit haemorrhagic disease virus (RHDV; single-stranded RNA) and myxoma virus (MYXV; double-stranded DNA) - were deliberately introduced to control wild rabbit populations in the 1950s and 1990s, respectively (Cooke & Fenner, 2002). Blowflies (Calliphoridae) and bushflies (Muscidae) are associated with the transmission of RHDV, while two species of rabbit fleas (*Spilopsyllus cuniculi* and *Xenopsylla cunicularis*) aid MYXV transmission (and mosquitos are potentially involved in the subsidiary transmission of both viruses) (Asgari, Hardy, Sinclair, & Cooke, 1998; Cooke & Fenner, 2002; Hall, Huang, Roberts, & Strive, 2019; McColl et al., 2002; Merchant et al., 2003; Sobey & Conolly, 1971). As viral replication is not believed to occur in insect tissue, transmission is entirely mechanical. RHDV is ingested by flies during feeding on carcasses and viable virus excreted in fly spots (Asgari et al., 1998), while fleas transmit MYXV through contaminated mouthparts (Fenner, Day, & Woodroofe, 1952). Although RHDV is transmissible directly by the faecal-oral route, flies facilitate transmission between isolated populations (Schwensow et al., 2014). Indeed, before it’s official release, RHDV escaped quarantine from Wardang island, South Australia, purportedly due to fly-vectored transmission (Asgari et al., 1998; McColl et al., 2002). MYXV can also be transmitted via direct contact, although biting insect vectors enhance transmission and as such, rabbit fleas were also deliberately introduced into Australia (Merchant et al., 2003; Sobey & Conolly, 1971).

Despite the importance of the ectoparasite-vector system in virus transmission and evolution, little is known about the composition of virus communities in both host types. Metagenomic studies of arthropod vector species such as mosquitoes and ticks have revealed an unexpectedly rich virus diversity, most of which likely do not infect vertebrates (Harvey, Rose, Eden, Lo, et al., 2019; Shi et al., 2017). Hence, it is not known what proportion of the viruses present in invertebrates pass to vertebrates and vice versa, although such information is central to understanding the evolution of vector-borne transmission and determining whether some viruses have more liberal host preferences than others.

The advent of bulk RNA sequencing (“meta-transcriptomics”) has revolutionized our perception of viral diversity and host range (Shi et al., 2016; Shi, Zhang, & Holmes, 2018), revealing large numbers of seemingly benign viruses (Shi, Lin, et al., 2018). The invertebrate meta-transcriptomic studies undertaken to date include various species of ectoparasite, such as mosquitos, ticks and fleas, revealing abundant and complex viromes (Harvey, Rose, Eden, Lawrence, et al., 2019; Harvey, Rose, Eden, Lo, et al., 2019; Shi et al., 2017). Herein, by comparing the viromes of Australian wild rabbits alongside associated rabbit fleas and sympatric flies, we present the first joint study of virome composition in vertebrates and their associated ectoparasites. Our aim was to determine whether and how virome composition differed between rabbits and the ectoparasites sampled on or near these rabbits, and whether some types of virus were common to both types of host such that they are involved in either biological or mechanical transmission.

## Materials and Methods

### Tissue Sampling

Sampling was performed at two sites within the Australian Capital Territory (ACT), Australia: site 1 was at CSIRO Crace (-35.22, 149.12), Gungahlin (GUN), a suburb of Canberra, while site 2 was at Gudgenby Valley (-35.74, 148.98) in Namadgi National Park (Gudg). At site 1, rabbits were trapped in carrot baited cages and killed by cervical dislocation. Trapping occurred over 3-5 consecutive nights for two separate weeks of the 2016/2017 southern hemisphere summer (18^th^ – 22^nd^ December 2016, 8^th^ – 11^th^ January 2017). A total of 20 rabbits were sampled, with weights ranging between 0.27 kg and 1.95 kg (mean 0.82 kg). At site 2, rabbits were killed by shooting on 2^nd^ February 2017. Eighteen rabbits were collected, weighing between 0.52 kg and 2.2 kg (mean 1.49 kg). Blood (in EDTA tubes), lung, liver, duodenum, and caecal content were collected from each rabbit. Where fleas were present on rabbits, they were collected and grouped by rabbit. Tissues and fleas were stored below -80°C immediately after collection.

Commercially available fly traps (Envirosafe^TM^) were placed at the same locations in the same weeks as rabbit sampling. Traps were baited with rabbit tissue/gut content and/or chicken necks, and bait was physically separated from flies to prevent contamination. Fly traps were left out for periods of up to 24 hours. Only live flies were taken from traps to ensure fresh samples. Live flies were chilled at 4°C or -20°C for periods of 5-10 mins to allow initial visual identification of fly species, before being frozen at -80°C. From site 1 (GUN), 149 flies representing 5 species were collected, while 22 flies from 2 species were collected from site 2 (Gudg) (Table 1).

**Table 1.** Rabbit and insect sampling and pooling details.

| Library | Site<sup>†</sup> | Species | Sample Type | No. samples in RNA-Seq pool |
| --- | --- | --- | --- | --- |
| **Rabbit tissues** |  |  |  |  |
| GUN-BI | 1 | *Oryctolagus cuniculus* | Blood | 20 |
| GUN-Li | 1 | *Oryctolagus cuniculus* | Liver | 20 |
| GUN-Lu | 1 | *Oryctolagus cuniculus* | Lung | 20 |
| GUN-Duo | 1 | *Oryctolagus cuniculus* | Duo | 20 |
| GUN-CC | 1 | *Oryctolagus cuniculus* | Caecal content | 20 |
| Gudg-BI | 2 | *Oryctolagus cuniculus* | Blood | 18 |
| Gudg-Li | 2 | *Oryctolagus cuniculus* | Liver | 18 |
| Gudg-Lu | 2 | *Oryctolagus cuniculus* | Lung | 18 |
| Gudg-Duo | 2 | *Oryctolagus cuniculus* | Duo | 18 |
| Gudg-CC | 2 | *Oryctolagus cuniculus* | Caecal content | 18 |
| **Arthropods** |  |  |  |  |
| GUN-F | 1 | *Spilopsyllus cuniculi*<sup>‡</sup> | Entire fleas grouped by rabbit | >70 fleas from 8 rabbits |
| GUN-ChV | 1 | *Chrysomya varipes* | Entire fly | 10 |
| GUN-Chsp | 1 | *Chrysomya rufifacies/albiceps*<sup>‡</sup> | Entire fly | 10 |
| GUN-CaV | 1 | *Calliphora vicina* | Entire fly | 10 |
| GUN-CaA | 1 | *Calliphora augur* | Entire fly | 10 |
| GUN-Sasp | 1 | *Sarcophaga impatiens*<sup>‡</sup> | Entire fly | 2 |
| Gudg-F | 2 | *Spilopsyllus cuniculi*<sup>‡</sup> | Entire fleas grouped by rabbit | >50 fleas from 7 rabbits |
| Gudg-CaA | 2 | *Calliphora augur* | Entire fly | 10 |
| Gudg-Un | 2 | *Musca vetustissima* | Entire fly | 2 |

<sup>†</sup>Site 1 - CSIRO Crace, Gungahlin; Site 2 - Gudgenby Valley in Namadgi National Park.
<sup>‡</sup>Species designation based on RNA-Seq data.

All work was carried out according to the Australian Code for the Care and Use of Animals for Scientific Purposes with approval from the institutional animal ethics committee (Permit CWLA-AEC#16-02).

### RNA extraction

RNA was extracted separately for each sample; from 20 mg of rabbit tissue or bone marrow, 75 µl of rabbit blood, individual whole flies or groups of at least 5 fleas from individual rabbits. RNA was extracted using the Maxwell 16 LEV simplyRNA tissue kit in combination with the Maxwell nucleic acid extraction robot (Promega, WI, USA), according to manufacturer’s instruction, including DNase treatment.

### Library construction and sequencing

Rabbit RNA was pooled by tissue type and collection site, with up to 20 individuals per pool, while insect RNA was pooled by species and collection site, with pool sizes ranging from 2 – 10 individuals (Table 1). Where large numbers of flies of the same species were collected, RNA from a maximum of 10 flies were pooled together. Liver RNA required further DNase treatment after pooling, using Invitrogen TURBO DNase (Thermofisher Scientific). All pooled RNA was further purified using the RNeasy MinElute clean-up kit (Qiagen, Hilden, Germany) and quantified using the Qubit RNA Broad-range Assay with the Qubit Fluorometer v3.0 (Thermofisher Scientific). RNA pools were assessed for quality using the Agilent RNA 6000 Nano kit and Agilent 2100 Bioanalyzer (Agilent Technologies, CA, USA). Library construction and sequencing was performed at the Australian Genomic Research Facility. Libraries were constructed using the TruSeq Total RNA Library Preparation protocol (Illumina, CA, USA) and rRNA was removed using the Illumina Ribo-Zero Gold rRNA removal kit (Epidemiology). Paired-end (100 bp) sequencing of each RNA library was performed on the HiSeq 2500 sequencing platform (Illumina, CA, USA).

### Assembly and genome annotation

*De novo* assembly of reads into contigs was performed using Trinity (Grabherr et al., 2011) following trimming with Trimmomatic (Bolger, Lohse, & Usadel, 2014). The RSEM tool (B. Li, Ruotti, Stewart, Thomson, & Dewey, 2010) in Trinity was used to calculate the relative abundance of each contig (expected counts). BLASTn and DIAMOND BLASTx were then used to compare Trinity contigs to the NCBI nucleotide (nt) database (*e*-value cut-off 1 x 10^-10^) and non-redundant protein (nr) database (*e*-value cut-off 1 x 10^-5^), respectively. Results were filtered so that only contigs that had a viral hit (excluding endogenous viruses/retroviruses) from each BLAST search were retained.

Equivalent BLAST analyses were performed on individual reads to detect viruses at low abundance, with *e*-value cut-offs of 1 x 10^-4^ for BLASTx and 1 x 10^-10^ for BLASTn. A conservative approach was taken such that only reads that had a virus result in both the BLASTn and BLASTx analyses were considered as legitimate hits. Ectoparasite library read-mapping to specific virus reference sequences or rabbit viral contigs was conducted using Bowtie2 (Langmead & Salzberg, 2012).

To remove residual host rRNA sequences, all reads were mapped to host rRNA using Bowtie2 (Langmead & Salzberg, 2012). The rabbit host rRNA target index was generated from a complete *O. cuniculus* 18S rRNA reference sequence obtained from GenBank (accession NR_033238) and a near complete *O. cuniculus* 28S rRNA sequence obtained from the SILVA high quality ribosomal database (Quast et al., 2013) (accession GBCA01000314). The arthropod rRNA target index was generated from 18S and 28S GenBank sequences from *Spilopsyllus cuniculi* and multiple *Chrysomya*, *Calliphora*, *Sarcophaga*, and *Musca* species. The total number of reads that did not map to host rRNA for each library were used as the denominator to calculate the percentage of reads mapped to viral contigs.

The Geneious assembler (Kearse et al., 2012) was used to extend viral contigs where possible. Open reading frames of viral contigs were identified using the online GeneMark heuristic approach to gene prediction tool (Besemer & Borodovsky, 1999), while conserved domains were identified using RSP-TBLASTN v2.6.0, a variant of PSI-BLAST (Altschul et al., 1997).

### Phylogenetic analyses

Reference RNA-dependent RNA polymerase (RdRp) amino acid sequences for each virus family were downloaded from NCBI and aligned with viral contigs using MAFFT v7.271 (Katoh & Standley, 2013). Where necessary, large data sets were condensed to a more manageable size using CD-HIT version 4.8.1 (W. Li & Godzik, 2006). Poorly and ambiguously aligned sites were removed using trimAl v1.2rev59 (Capella-Gutierrez, Silla-Martinez, & Gabaldon, 2009). Alignments were visualized in Geneious (Kearse et al., 2012). Maximum likelihood trees of each alignment were inferred using PhyML (Guindon et al., 2010) employing the LG amino acid replacement model selected by IQTree (Nguyen, Schmidt, von Haeseler, & Minh, 2015), using a combination of NNI (Nearest Neighbour Interchange) and SPR (Subtree Pruning and Regrafting) branch-swapping. Branch supports were estimated with the Shimodaira-Hasegawa (SH)-like approximate likelihood ratio test (Guindon et al., 2010). The size and length of each alignment is provided in Table S1 and details of viral contigs included in phylogenies are provided in Table S2.

### Screening PCRs for detection of rabbit calicivirus and picornavirus

Primer sets were designed to amplify a small region of each of the novel rabbit calicivirus and picornavirus genomes for the detection of these viruses in individual caecal content samples. The calicivirus primer set GRC_F5.6 (5’-TTA CTC AGA GCG ACC AAG TGC-3’, positive sense) and GRC_R5.9 (5’-CCA GTT CTC GCC TGT ATC CAG-3’, negative sense) amplified a 278 bp region, while the picornavirus primer set GRP_F6.5 (5’-GAT CTT ATC CCA CCC AAT CGT GA-3’, positive sense) and GRP_R6.9 (5’-ATA GCC TCT TCT CCA TAA CCA AGC-3’, negative sense) amplified a 401 bp region. RT-PCRs were conducted using the QIAGEN® OneStep Ahead RT-PCR Kit according to the manufacturer’s directions with 1 µl of RNA (diluted 1:10 in nuclease-free water) in a 10 µl reaction volume with 0.25 µM of each primer. PCR conditions included 10 cycles of touchdown PCR, with the annealing temperature decreasing by 0.5 °C each cycle from a starting temp of 60 °C, and a further 30 cycles with annealing temperature at 55 °C. Representative amplicons were Sanger sequenced to confirm their legitimacy.

### Extension/confirmation of 3’ end of novel calicivirus genome

First strand cDNA synthesis was conducted using the Invitrogen Superscript IV reverse transcriptase system (Thermofisher Scientific, MA, USA), with 5 µl of RNA and 0.5 µM of GV270 gene-specific primer (Eden, Tanaka, Boni, Rawlinson, & White, 2013) in a 20 µl reaction volume. PCR was conducted using the Invitrogen Platinum Taq Polymerase High Fidelity kit according to the manufacturer’s protocol with specifically designed forward primer GRC_F6.2 (5’-CAG AGA ATG AGC TCA ACC GAC A-3’), and reverse primer GV271 (Eden et al., 2013). Reaction volumes of 40 µl included 2.5 µl of cDNA template and 1 µM of each primer. PCR was conducted for 45 cycles, with the annealing temperature starting at 65 °C and decreasing by 0.5 °C each cycle. The positive amplicon was approximately 500 bp (includes polyA tail) and was Sanger sequenced for confirmation.

### Detection of lagoviruses in flies and rabbit carcasses

RNAs from individual flies and from the bone marrow of rabbit carcasses found near Gungahlin fly traps were screened for the presence of pathogenic lagoviruses using the multiplex RT-PCR described previously (Hall et al., 2018).

## Results

### Genetic identification of unknown arthropods

The majority of arthropods analysed in this study were identified to the species level through visual inspection. The remainder were characterised using the RNA-Seq data. Fleas were confirmed to be *Spilopsyllus cuniculi* (rabbit fleas) based on the presence of several highly abundant contigs of *Spilopsyllus cuniculi* rRNA and EF1a genes and the absence of any other *Spilopsyllus* species genes. A library of unidentified *Chrysomya* species (GUNChsp library) was determined to be *Chrysomya rufifacies* or *albiceps* (these two species are potentially the same) based on EF1a and rRNA genes. An unknown *Sarcophaga* species was most likely *Sarcophaga impatiens* based 28S rRNA identity.

### Fly species trapped

A wider diversity of flies were trapped at site 1, a suburb of Canberra (n = 5 species), than at site 2, in Namadgi National Park (n = 2 species, Table 1). Species from the genera *Calliphora*, *Chrysomya* (both Calliphoridae) and *Sarcophaga* (Sarcophagidae) were collected from site 1, while species from *Calliphora* and *Musca* (Muscidae) were isolated from site 2. *Calliphora augur* was the only species trapped at both sites (Table 1).

### Virus contigs in ectoparasites

A large number of RNA viral contigs were assembled from the flea and fly libraries. Of the invertebrate species, *Calliphora vicina* had the highest virus abundance (as a proportion of non-rRNA reads), with almost 2% of non-rRNA reads being viral, while *Chrysomya* species had viral abundances of only 0.013 – 0.019% (Figure 1). Each ectoparasite species had virus contigs from between 6 and 14 different RNA virus families. While viruses from several different families were detected in fleas (10 – 14 families), viruses in both flea libraries largely belonged to the *Iflaviridae* and Sobemo-like viruses (Figure 1). Of the fly species, the *Calliphora augur* libraries harboured the highest number of virus families, although diversity did not differ extensively between libraries. Although only fleas and *Calliphora augur* were sampled from both sites, the viral diversity of these two species at each site suggests that viral composition was associated with host species rather than collection location (Figure 1). The fly results also suggest that there is a trend in viral composition at the genus level (GUNCaA, GudgCaA and GUNCaV are genus *Calliphora*, while GUNChsp and GUNChV are *Chrysomya*), with decreasing similarity in viral composition at the family level and beyond (Figure 1). Members of the *Partitiviridae* and *Phenuiviridae* were detected in all invertebrate species, although it is possible that some of the low abundance viruses (as low as 46 and 78 reads, respectively) represent cross-contamination since all flies were caught in the same trap at each location.

**Figure 1.**
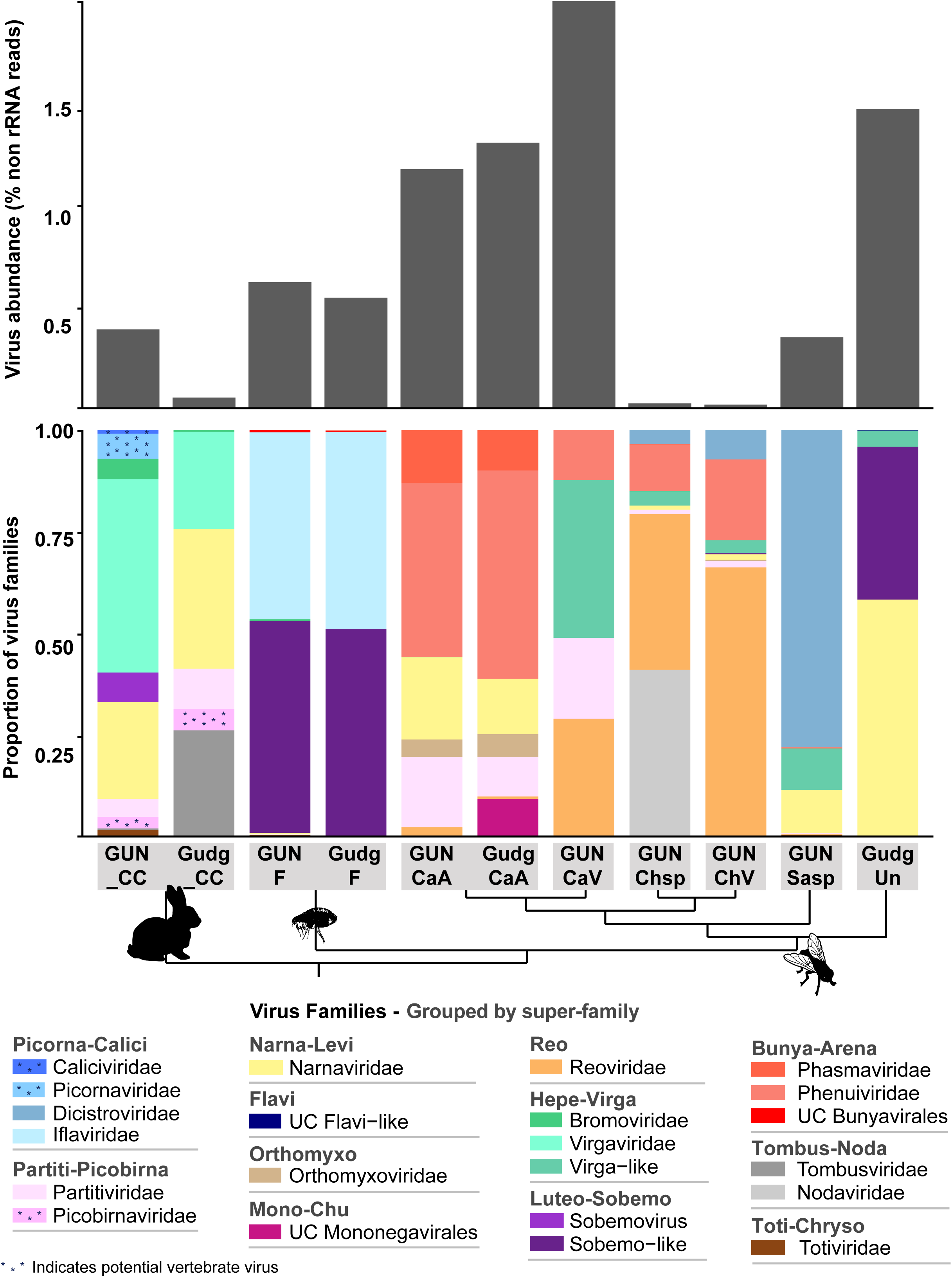
RNA virus abundance and composition in rabbit and invertebrate libraries. The top plot displays the abundance of viral reads (y-axis) in each library (x-axis) as a proportion of total non-rRNA reads. The bottom plot shows the viral composition of each library by virus family/group (shaded/grouped by superfamily). Potential vertebrate viruses are indicated by asterisks within the shading. Only RNA viruses (with RdRp) are shown and only virus families that had an abundance of at least 0.001% in at least one library are presented. UC denotes unclassified viruses. Virus libraries are labelled as follows (collection site - species): GUN_CC, Gungahlin - rabbit caecal content; Gudg_CC, Gudgenby - rabbit caecal content; GUNF, Gungahlin - flea; GudgF, Gudgenby - flea; GUNCaA, Gungahlin - *Calliphora augur*; GudgCaA, Gudgenby - *Calliphora augur*; GUNChsp, Gungahlin - *Chrysomya rufifacies/albiceps*; GUNCaV, Gungahlin - *Calliphora vicina*; GUNChV, Gungahlin - *Chrysomya varipes*; GUNSasp, Gungahlin - *Sarcophaga impatiens*; GudgUn, Gudgenby - *Musca vetustissima*. Note that only the caecal content libraries from rabbits are included in the plots since no viruses were found in the other libraries. A cladogram connecting the libraries beneath the x-axis indicates the relationships between the sampled hosts in each library, where tips represent host species and nodes from top-to-bottom represent the levels of genus, family, super-family, order, class and kingdom.

To establish the diversity and potential host of newly defined viruses, family level (in some cases super-family level) phylogenetic trees were estimated using the virus RdRp (Figure 2). While many of the highly diverse phylogenies had poorly resolved topologies, we identified at least 25 diverse viruses that likely constitute new species. The majority of viruses found in invertebrate species clustered with invertebrate-associated viruses in the *Dicistroviridae, Iflaviridae, Nodaviridae,* Flavi-like, *Solemoviridae/*Sobemo-like, Virga-like*, Orthomyxoviridae*, *Mononegavirales, Reoviridae, Phasmaviridae (Bunyavirales)* and unclassified bunyavirales groups. Additionally, many of the viruses found in insects, particularly fleas, were potentially viruses of fungi, protozoa or algae, being present in the *Hypoviridae, Narnaviridae, Partitiviridae,* the *Totiviridae-Chrysoviridae* group and certain *Phenuiviridae (Bunyavirales)*. The *Bromoviridae* virus identified in Gungahlin fleas clusters firmly among plant viruses, and with an abundance of only 0.002% it likely represents a plant virus carried by fleas rather than a virus of fleas themselves. Indeed, care must be taken in assigning viruses to hosts on the basis of metagenomic data alone. The *Iflaviridae* flea viruses found in this study clustered most closely with Watson virus, detected in fleas (*Pygiopsylla*) from Australian marsupials (Figure 2).

**Figure 2.**
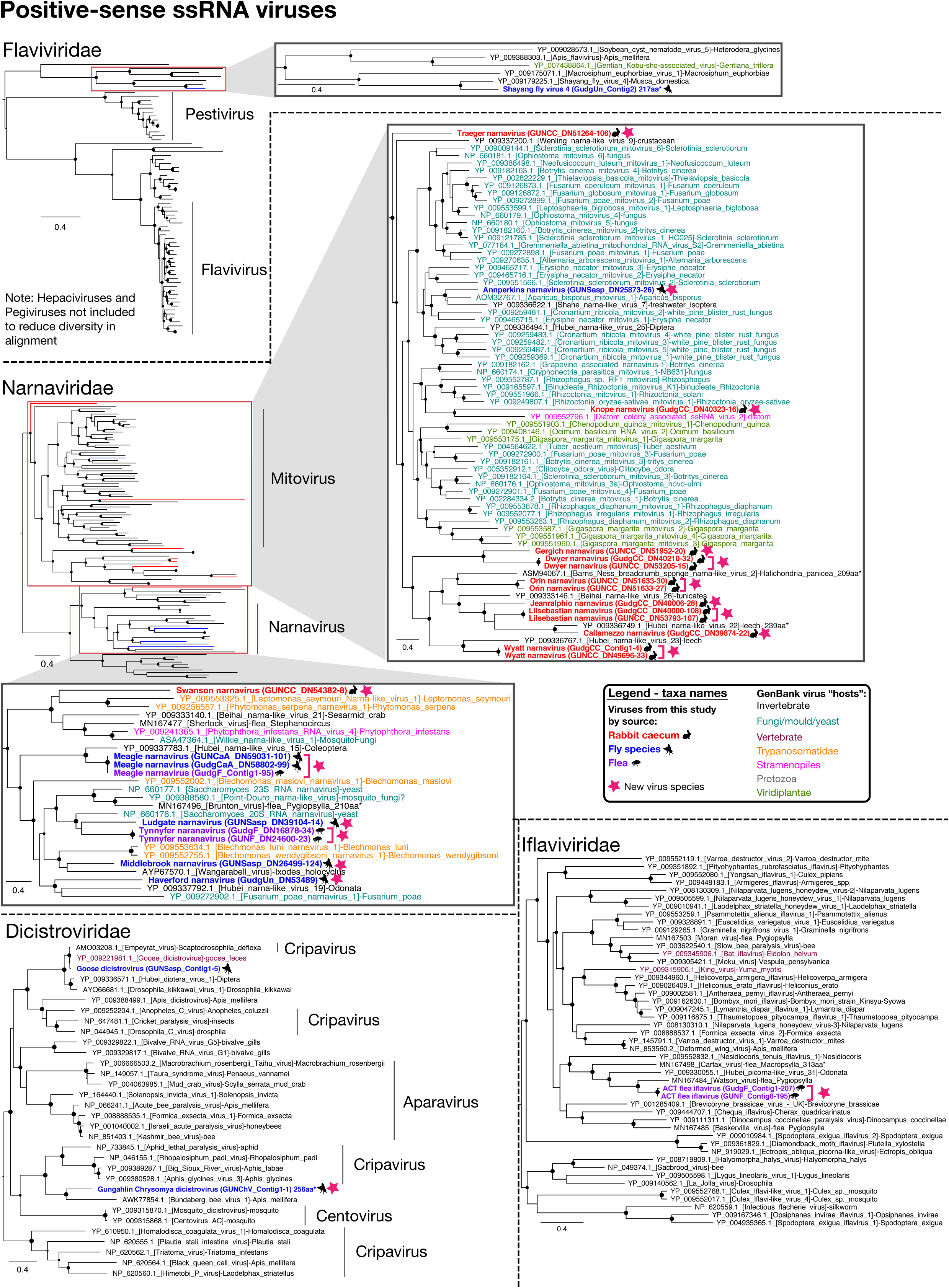
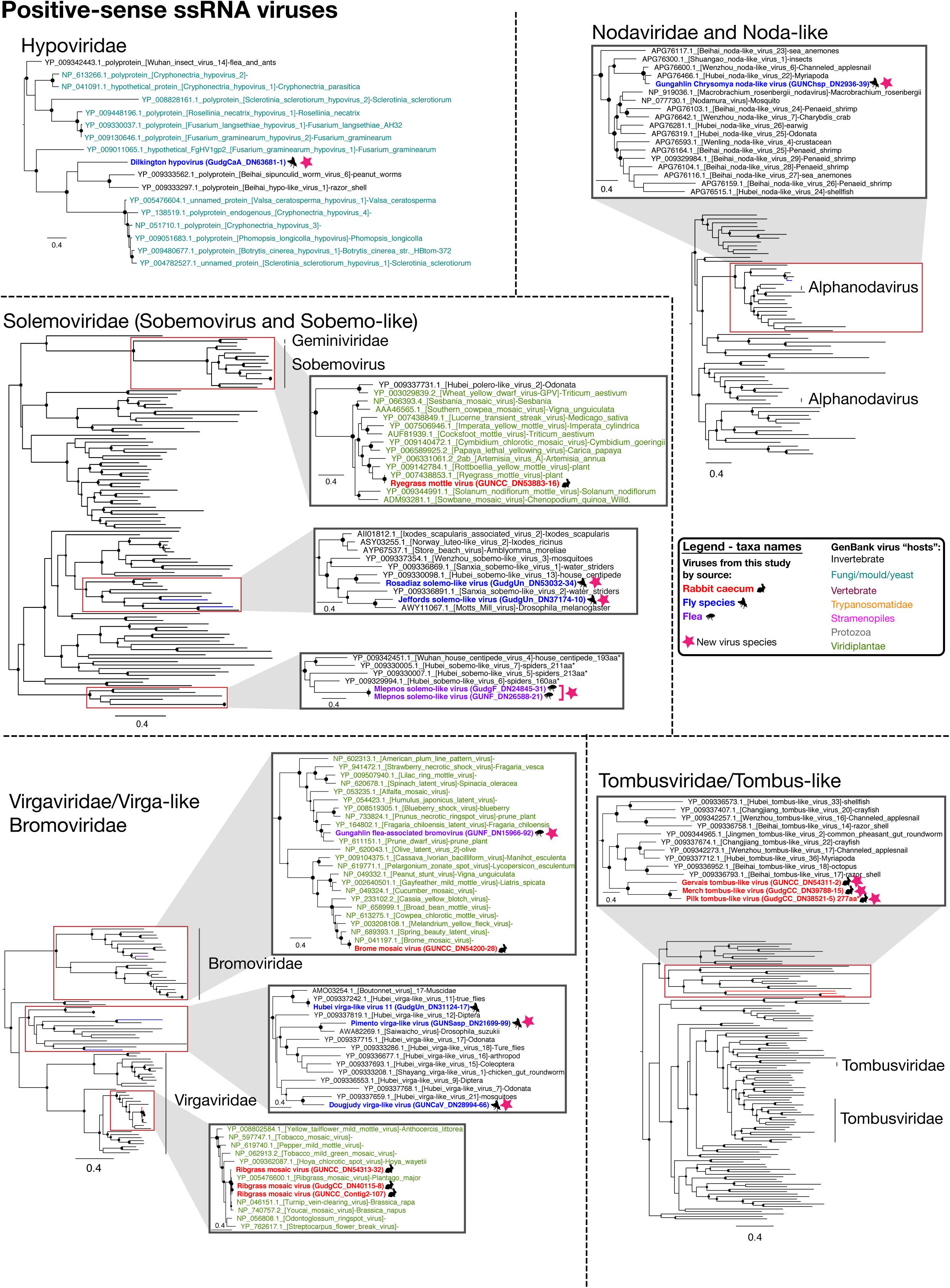
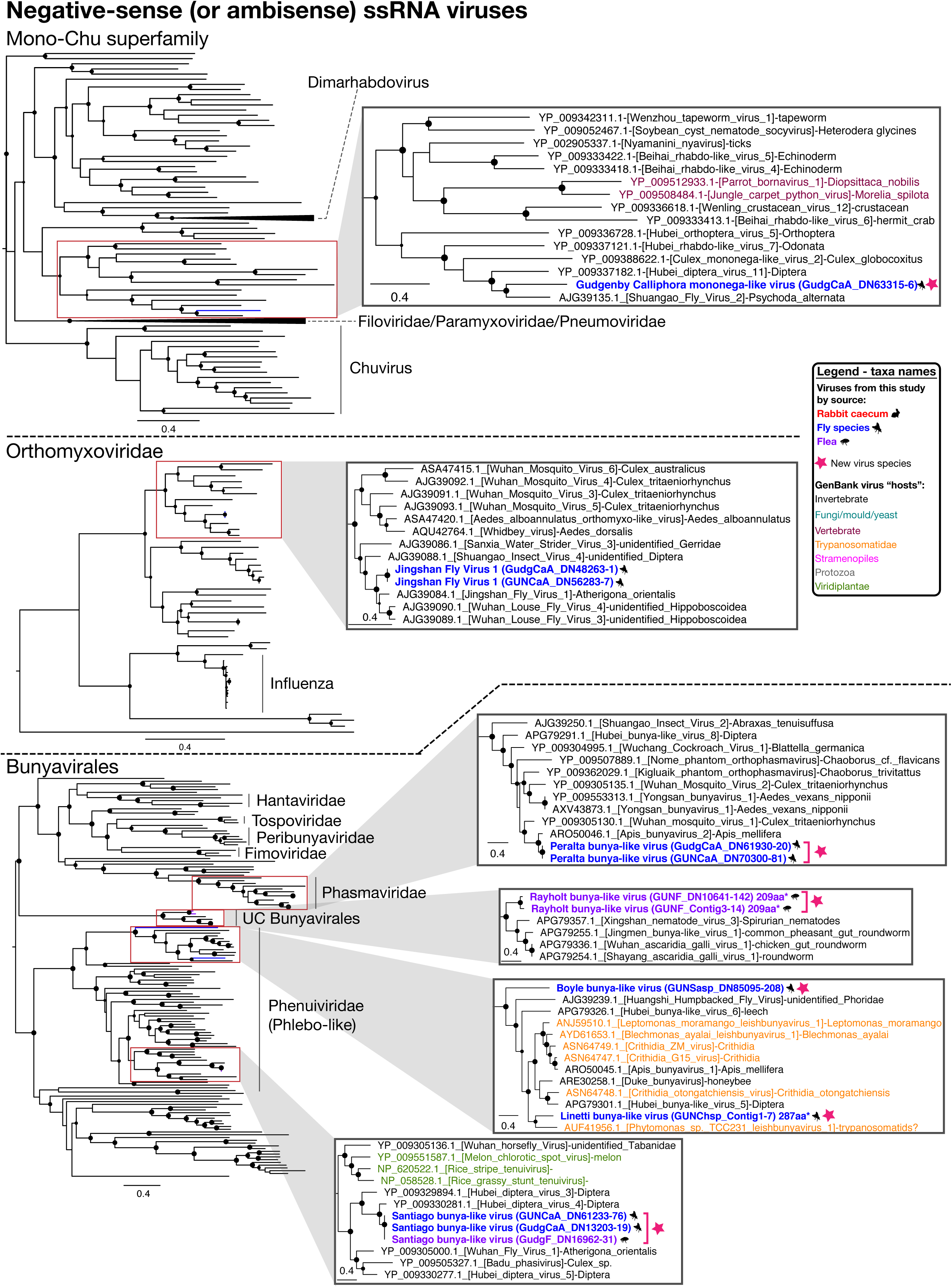
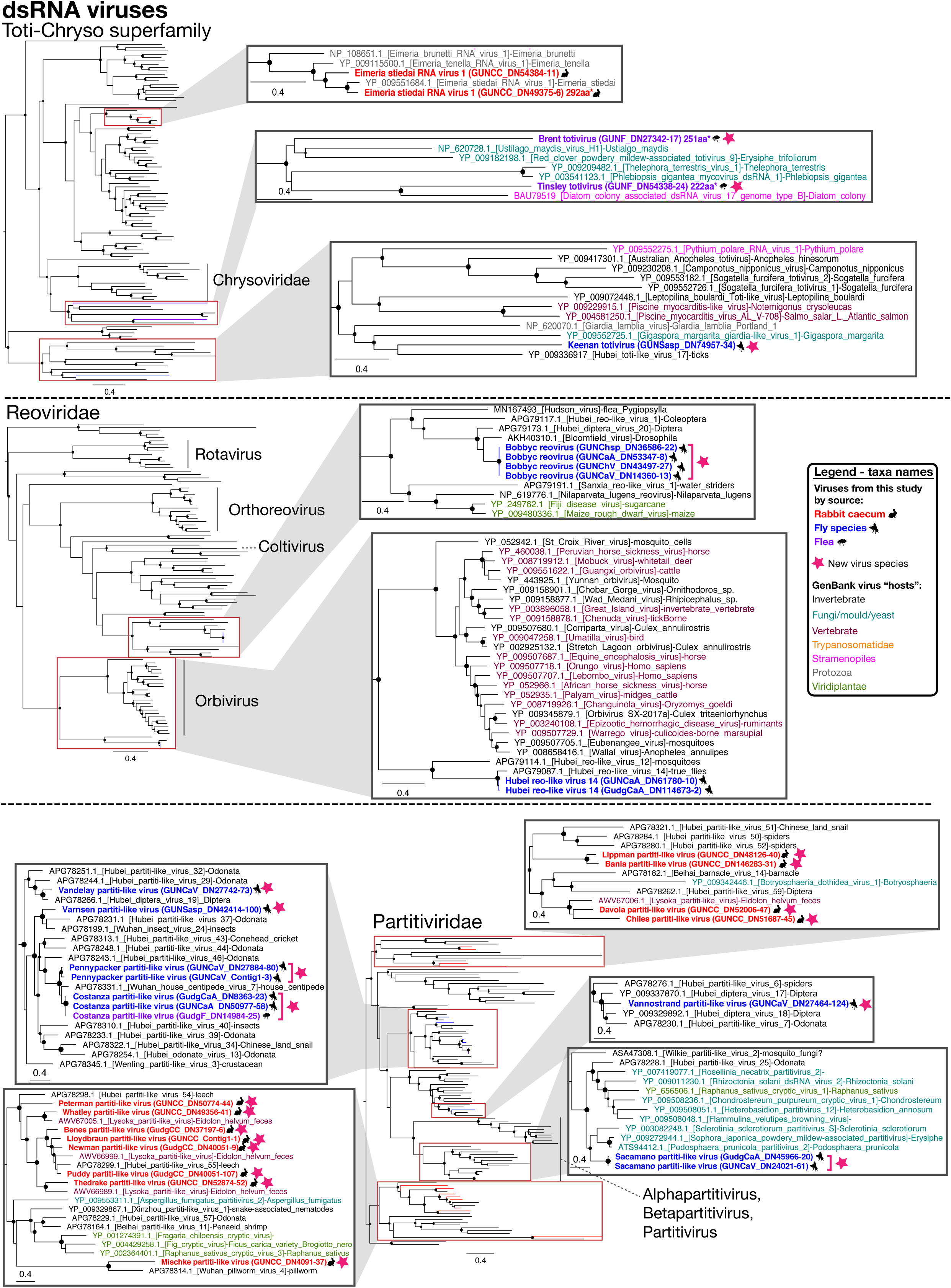
ML trees of the RdRp of likely non-vertebrate viruses. The taxa name (and branches in minimized trees) for sequences obtained in this study are bolded and coloured red (rabbit caecal content), blue (flies), or purple (fleas), based on the animal from which they were obtained, with relevant animal symbols adjacent to the names. Viruses that likely constitute a new viral species are indicated by a pink star symbol adjacent to taxa names, and a proposed virus species name is given as the taxa name (with strain name in parentheses). For GenBank sequences, taxa names are coloured by the apparent host group from which virus or viral sequence was reportedly isolated: black, invertebrate; teal, fungi/mould/yeast; maroon, vertebrates; orange, Trypanosomatidae; pink, Stramenopiles (microalgae(diatom)/Oomycetes); grey, other protozoa (Coccidia, *Trichomonas*, *Giardia*). SH-like support values are represented by circles at the nodes if >0.7 and are sized according to values where the largest circles represent an SH-like support of 1. For sequences that are less than 80% of the alignment length, the sequence length in amino acids (aa) and an asterisk is included in the taxa name.

### Virus contigs in rabbits

No viral contigs could be assembled from the rabbit liver, duodenum, or lung libraries. A small number of viral contigs were found in the Gudgenby blood library, but these were potential contaminants since (i) rabbits from Gudgenby were shot, occasionally resulting in perforation of the caecum which would contaminate blood in the body cavity, (ii) all viruses detected in the blood were also detected in the caecum (including plant viruses unlikely to be in blood), and (iii) no viruses were found in the blood of rabbits from the Gungahlin site where there was no body cavity contamination.

In contrast, the caecal content for rabbits from both sites contained many viruses, with 8 and 11 RNA viral families detected in the Gudgenby and Gungahlin rabbits, respectively (Figure 1), including over 25 likely new viral species. The viral composition of rabbit caecal content was less consistent between the two sites than for the invertebrates. This may be a consequence of sampling only small sections of caecal content, but could also reflect differences in diet at each site (predominantly introduced pastures at site 1 versus more subalpine native grassland plants at site 2). Regardless, *Narnaviridae* and *Virgaviridae* were both highly abundant in the caecal content of rabbits from both locations, while *Tombusviridae* was also a major component of the caecal virome of Gudgenby rabbits (Figure 1). These three virus families, that make up more than 70% of the total viral abundance in rabbit caecal content at each site, likely represent viruses of the rabbit diet (plants) and commensal/parasitic organisms such as fungi and protists. Although the *Tombusviridae* were traditionally associated with plants, recent studies have found many tombus-like viruses in invertebrates (Shi et al., 2016) and these group with the caecal content viruses determined here. Hence, these tombus-like viruses may in fact infect commensal or parasitic microorganisms such as protists or fungi, or rabbits may be incidentally eating invertebrates.

Although less abundant, diverse novel viruses from two vertebrate viral families - the *Caliciviridae* and the *Picornaviridae* - and one potentially vertebrate-associated viral family, the *Picobirnaviridae,* were detected in rabbit caecal content at both sites: all three at Gungahlin, and the *Picobirnaviridae* at Gudgenby. Two related *Caliciviridae* contigs were assembled, with 77.8% nucleotide identity in the genome and 90.8% identity at the RdRp protein level. They clustered most closely - although distantly - with a pig calicivirus and marmot norovirus (Figure 3), sharing 52 - 54% identity in the RdRp. Such a divergent phylogenetic position suggests that the new calicivirus contigs represents a new viral species (Figure 3), which we have termed *Racaecavirus*. After Sanger sequencing to extend the 3’ end, one of the racaecavirus contigs encompassed a near complete genome, missing only the 5’ UTR. *Racaecavirus* exhibited a classic calicivirus-like genome organization, with two open reading frames (ORF), one encoding a polyprotein including RdRp and capsid domains, and the second encoding a small protein of unknown function (Figure 3). Oddly, there appears to be only 1 nucleotide in the 3’ UTR of this genome sequence, which was confirmed by Sanger sequencing.

**Figure 3.**
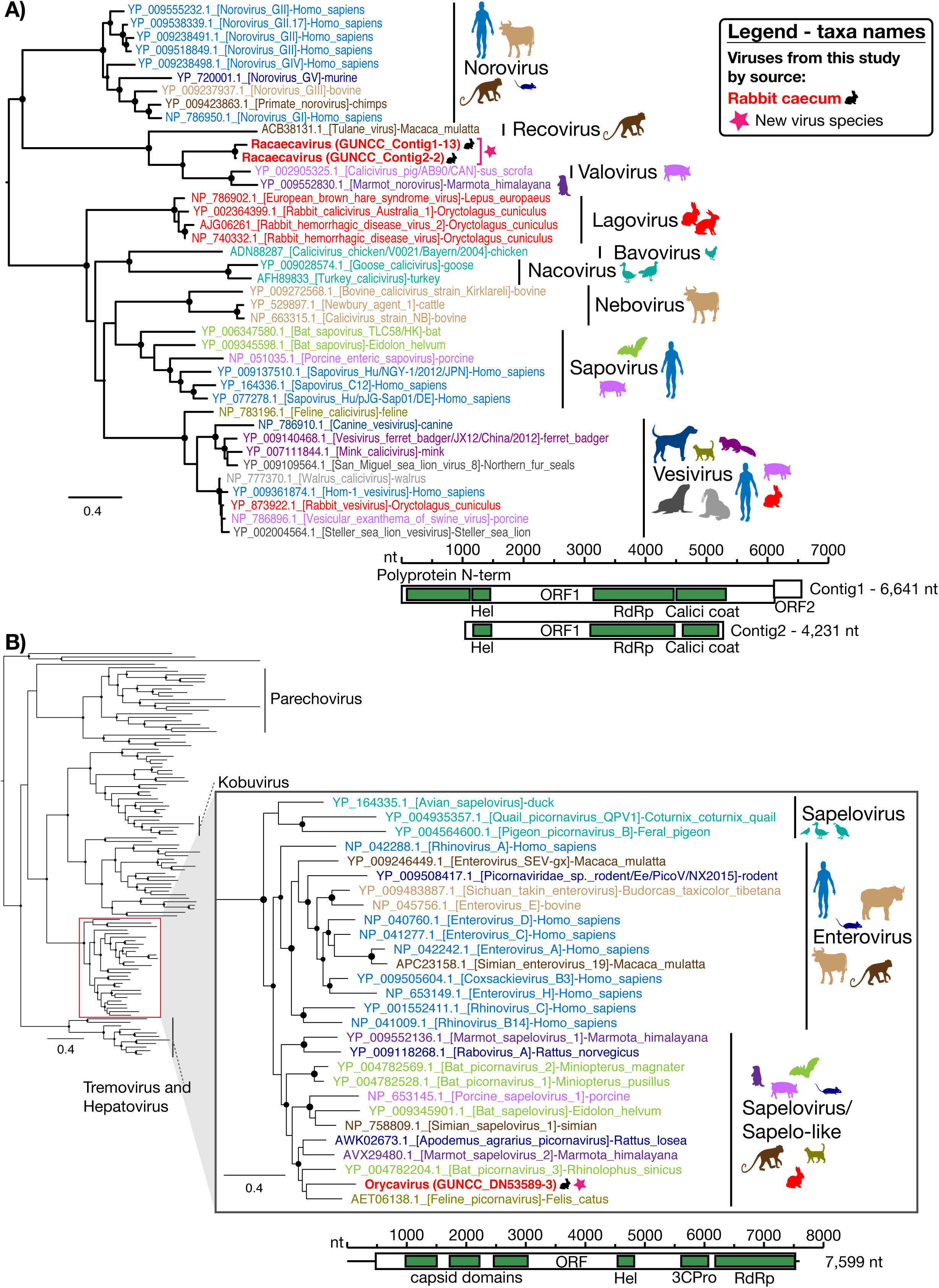
Phylogenetic analysis of the RdRp of vertebrate-specific viruses found in rabbit caecum. ML trees of the RdRp region of (A) the novel rabbit calicivirus – *Racaecavirus* - and (B) the novel rabbit picornavirus – *Orycavirus* - together with representative reference sequences for these virus families are shown. Taxa names of the viruses discovered in this study are bolded with a black rabbit symbol adjacent. A pink star symbol adjacent to taxa names indicates a novel virus species and the proposed virus species name is given as the taxa name (with strain name in parentheses). GenBank accession numbers are included in the taxa name and these names are colour-coded according to host as specified by coloured symbols to the right of each tree. Clade labelling indicates specific genera. SH-like approximate likelihood ratio branch support greater than 0.7 is indicated by circles at the nodes which are sized according to degree of support (SH-like support of 1 has the largest size). Trees were midpoint rooted for clarity. The genome structure and length of the isolated contigs is shown below each tree, with open boxes representing ORFs, and green boxes indicating conserved protein domains: Polyprotein N-term, N-terminal region of the polyprotein; Hel, helicase; RdRp, RNA-dependent RNA polymerase; Calici coat, calicivirus capsid/coat protein; 3CPro, 3C proteinase.

Similarly, the entire coding region was obtained for a novel member of the *Picornaviridae*. This contained one large ORF, typical of the *Picornaviridae*, with multiple capsid proteins preceding non-structural proteins (Figure 3). The sequence also contained a 5’ (478 nt) and 3’ (74 nt) UTR, although it is not clear if these are complete. The novel virus clusters, with strong support, with members of the *Enterovirus* and *Sapelovirus* genera (Figure 3). The closest sequenced relatives were feline picornavirus, bat picornavirus 3, Apodemus agrarius picornavirus, and marmot sapelovirus 2, that share an identity of 61 - 64% with the novel rabbit picornavirus in the RdRp protein (Figure 3). This level of divergence and phylogenetic position would define this virus as a new species within the genus *Sapelovirus* or a newly defined sapelovirus-like genus (Figure 3) (Zell, 2018). Accordingly, we propose the name *Orycavirus*.

Since RNA-sequencing was conducted on pools of 18-20 samples, specific RT-PCRs for the novel racaecavirus and orycavirus identified here were designed to determine their frequency in individual animals. As these two viruses were only found in the Gungahlin caecal content library, only Gungahlin samples were screened. Racaecavirus was detected in 4/20 samples, while orycavirus was detected in 10 of the 20 samples tested.

Finally, several picobirnaviruses were identified in rabbit caecal content, all of which clustered strongly in the supposedly vertebrate-specific genogroup 1 clade (Figure 4). Based on individual species sharing <75% amino acid similarity in the RdRp alignment, these data likely contain nine novel picobirnaviruses (although defined species demarcation criteria for this family are lacking). Consistent with naming conventions adopted for most picobirnavirus species, the tentative new viruses were named *Rabbit picobirnavirus* 1-9. Importantly, these viruses did not form a monophyletic group, but were distributed throughout genogroup 1. This pattern is typical of the *Picobirnaviridae* that show limited host structure in the RdRp phylogeny (Figure 4), and is compatible with the idea that these are in fact bacterial-associated viruses (Krishnamurthy & Wang, 2018). The RdRp segments (segment 2) were predicted to have one ORF, consistent with other members of this family. While pairing segments was difficult, several longer picobirnavirus segments with at least one large ORF, likely encoding the capsid, were identified in both caecal content libraries.

**Figure 4.**
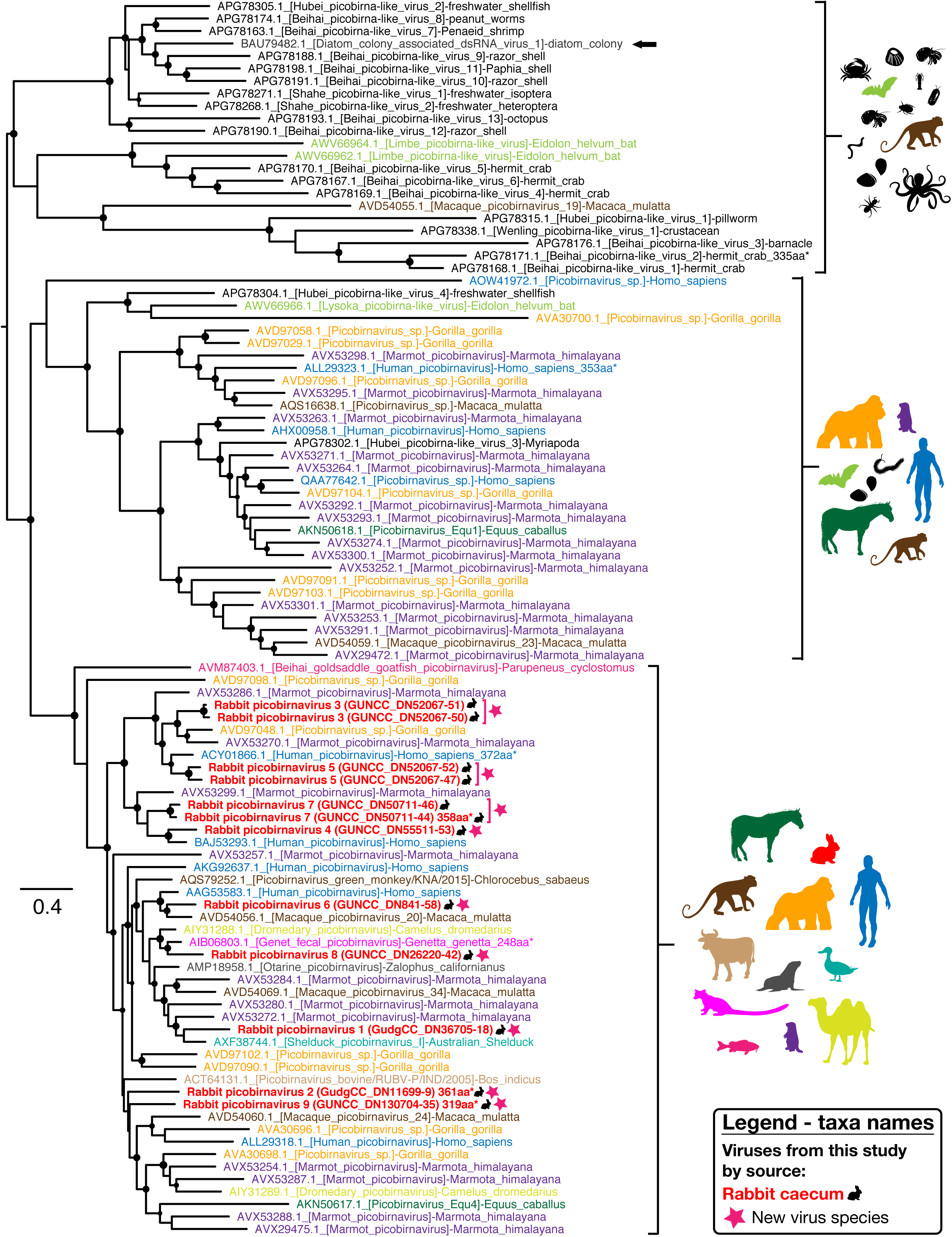
Phylogenetic analysis of the RdRp of novel picobirnaviruses. ML tree of the RdRp region of novel rabbit picobirnaviruses and representative picobirnaviruses from GenBank. The novel rabbit picobirnavirus taxa names are bolded, coloured red and emphasised with a black rabbit symbol adjacent to the name. A pink star symbol adjacent to taxa names indicates a novel virus species and the proposed virus species name is given as the taxa name (with strain name in parentheses). The taxa names of GenBank sequences include accession numbers and are coloured according to the host taxa from with they were isolated (all invertebrate host taxa are coloured black). The host taxa associated with sequences in each clade are indicated with symbols to the right of the clade. The single picobirnavirus sequence isolated from a diatom colony is indicated with an arrow. SH-like approximate likelihood ratio branch support greater than 0.7 is indicated by circles at the nodes which are sized according to degree of support (SH-like support of 1 is maximum size). Trees were midpoint rooted for clarity.

### Virus families present in both insect and rabbit libraries

Viral contigs from the *Virgaviridae/Bromoviridae*/Virga-like (plant/invertebrate-associated), *Solemoviridae*/Sobemo-like (plant/invertebrate-associated), *Narnaviridae* (fungi/parasites/invertebrate-associated), *Partitiviridae* (plant/invertebrate/fungi/vertebrate faeces-associated), *Tombusviridae* (plant/invertebrate-associated) and *Toti-Chryso* (parasites/invertebrates/fungi-associated) groups were assembled from rabbit caecal content as well as from both fleas and flies (Figure 1 and Figure 5). As noted above, the viruses assembled from rabbit caecal content in these virus families were unlikely to be actively replicating in rabbits. In addition, where viruses of the same family were assembled from arthropods as well as rabbits, they did not cluster together (Figure 2).

**Figure 5.**
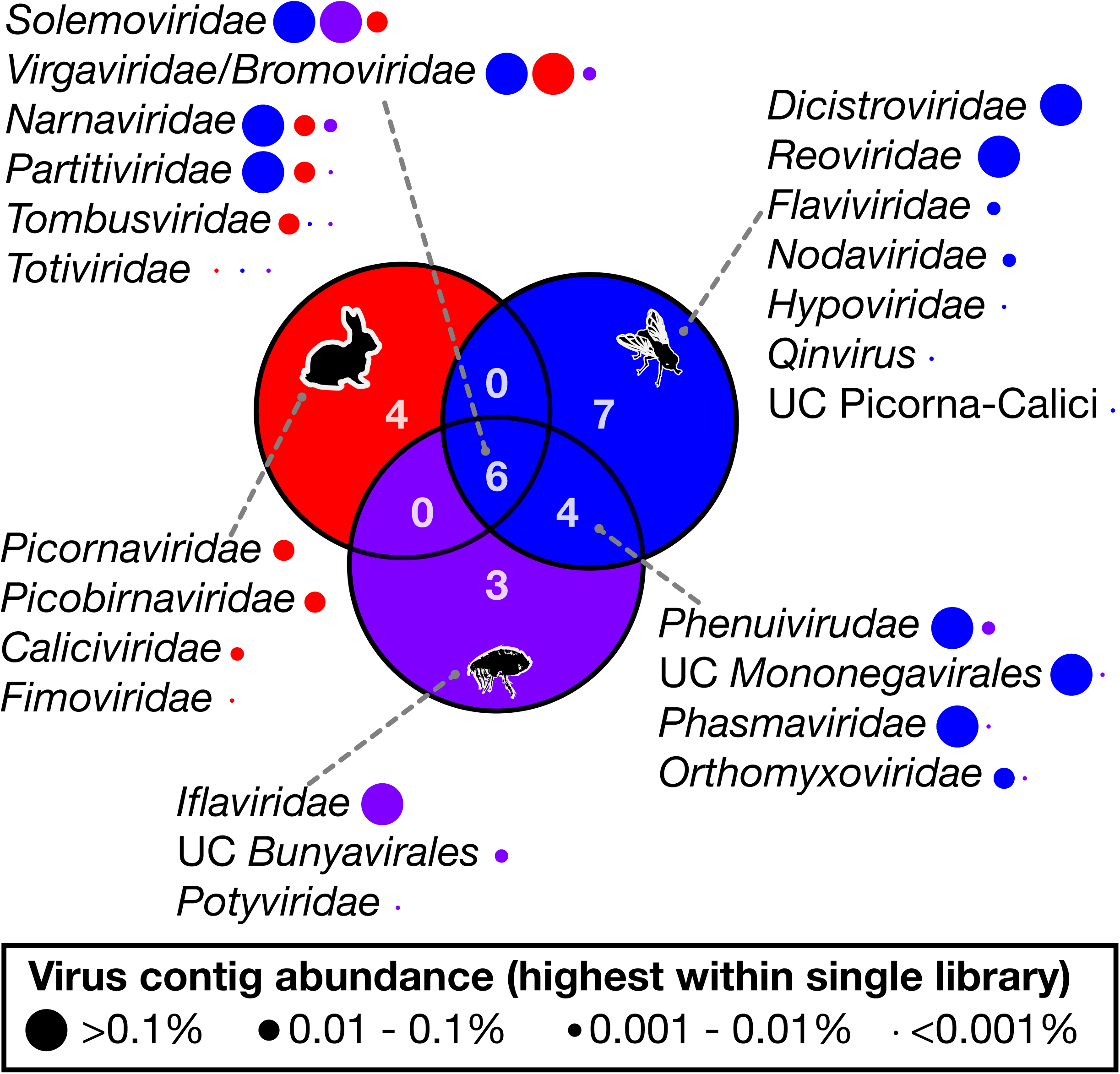
Overlap of RNA viral families in rabbits and ectoparasites. The number of viral families/groups for which contigs were assembled from either rabbit caecal content libraries (red circle), fly libraries (blue circle) and flea libraries (purple circle), and the level of overlap for each host group are indicated by a Venn diagram. The viral families associated with each segment are listed with grey dotted lines connecting lists to segments of the Venn diagram. The abundance of each viral family for the three groups is indicated by the size of circles next to virus family names. Circles are colour-coded according to the rabbit, fly or flea group with which they are associated, and the circle sizes reflect the highest abundance of the relevant virus family within a single library in the rabbit, flea or fly group. UC=unclassified.

To further investigate the viral overlap between rabbits and ectoparasites, reads from ectoparasite libraries were mapped to the viral contigs from the rabbit caeca. A total of 58 viral reads mapped to rabbit virus contigs, all associated with the viral groups described above, and hence were likely mapping to conserved regions. Taken together, these results show that no abundant viral species were shared between host and ectoparasites, such that there was no strong evidence of biological vector transmission.

### Low abundance vertebrate-associated viruses in ectoparasite libraries

If the ectoparasites studied here were involved in mechanical transmission, viruses may not be sufficiently abundant to be assembled into contigs. To detect vertebrate viruses at low abundance we subjected individual reads from the flea and fly libraries to BLASTn and BLASTx analyses. Accordingly, small numbers of reads were detected for two known rabbit-specific viruses (Figure 6): Lagoviruses (RHDV and related viruses) of the *Caliciviridae* family and rabbit astroviruses. The lagovirus reads detected included RHDV, rabbit haemorrhagic disease virus 2 (RHDV2), and the benign rabbit calicivirus Australia-1 (RCV-A1). Because of recombination between RHDV, RHDV2, and RCV-A1 (Hall et al., 2018), classification of these viruses based on small numbers of reads is difficult. However, the presence of reads mapping to the non-structural gene segments of RHDV and the RCV-A1-like viruses, as well as the structural gene segments of RHDV2, suggests the presence of at least two RHDV-like viruses in these fly libraries - a recombinant RHDV/RHDV2 and recombinant RCV-A1-like/RHDV2. Equivalent read BLAST analyses were conducted on rabbit libraries: two reads from RHDV2 recombinants were found in each of the Gudgenby liver, Gudgenby lung and Gungahlin blood libraries. Since they were at very low abundance, these viruses were not likely to be actively replicating in these rabbits, although they may represent the early hours of infection or a cleared infection. No vertebrate-specific virus reads were detected in the flea libraries or Gudgenby fly libraries. Due to the difficulty in confirming the legitimacy of viral reads, only those that had a virus result for both BLASTx and BLASTn analyses were included. Hence, this method will have necessarily led to a conservative estimate and the omission of diverse virus reads since these are unlikely to be detected in a BLASTn analysis.

**Figure 6.**
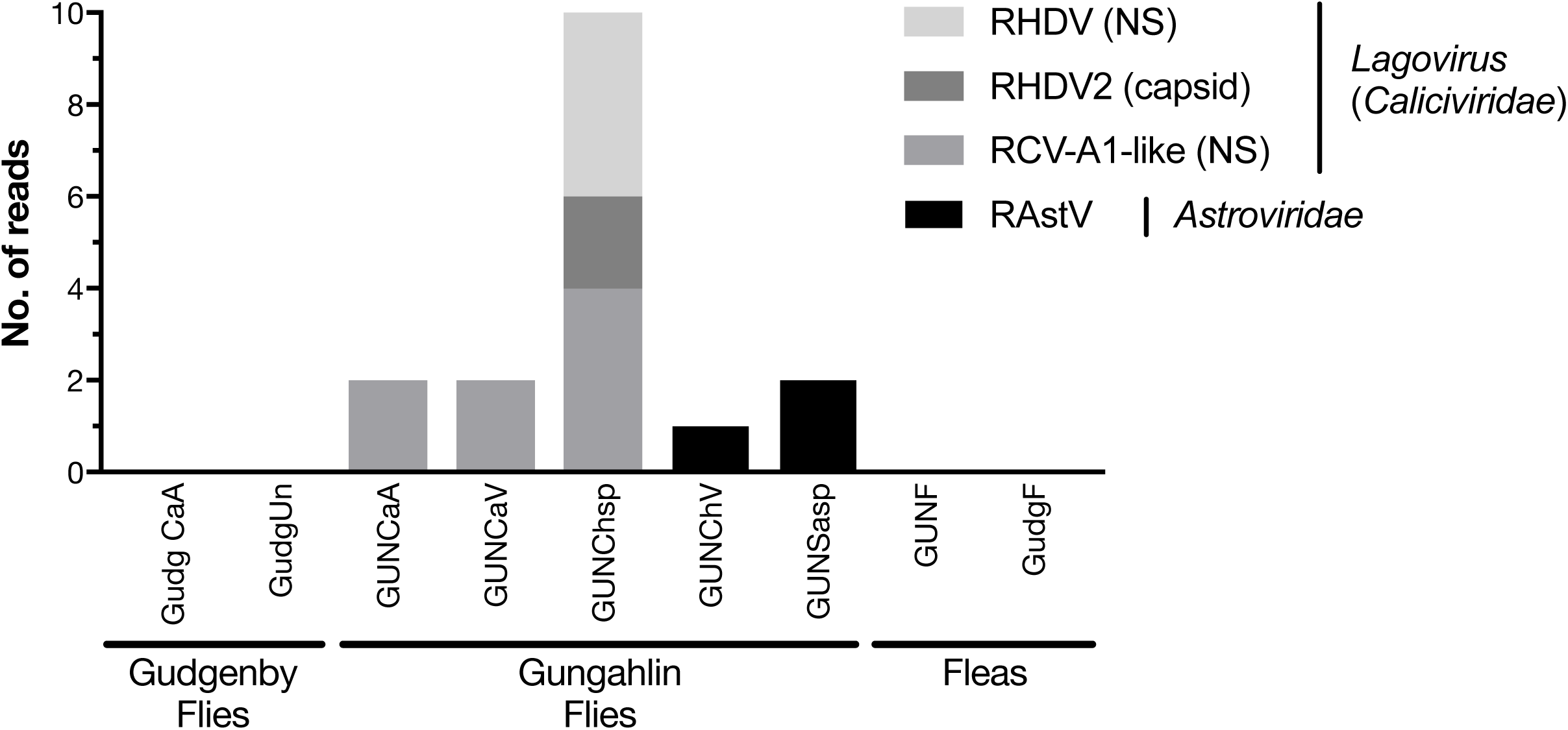
Vertebrate-specific virus reads detected in ectoparasite libraries. The number of reads from vertebrate viruses (y-axis) detected in ectoparasite libraries (x-axis) is presented as a stacked bar plot. Vertebrate virus reads that were detected include rabbit astrovirus (RAstV), and RHDV-like viruses that include recombinant variants of rabbit haemorrhagic disease virus (RHDV), and related viruses rabbit haemorrhagic disease virus 2 (RHDV2), and rabbit calicivirus Australia 1 (RCV-A1). In the legend, NS and capsid in parentheses indicates reads mapping to non-structural genes or capsid gene, respectively. Ectoparasite libraries are labelled as follows (location - species): GUNF, Gungahlin - flea; GudgF, Gudgenby - flea; GUNCaA, Gungahlin - *Calliphora augur*; GudgCaA, Gudgenby - *Calliphora augur*; GUNChsp, Gungahlin - *Chrysomya rufifacies/albiceps*; GUNCaV, Gungahlin - *Calliphora vicina*; GUNChV, Gungahlin - *Chrysomya varipes*; GUNSasp, Gungahlin - *Sarcophaga impatiens*; GudgUn, Gudgenby - *Musca vetustissima*.

Since some viruses were represented by as little as a single read per library, we confirmed the presence of RHDV-like viruses in invertebrates by RT-PCR. Importantly, several individual flies from all three libraries with RHDV-like reads were positive by RT-PCR despite each library having only 2 – 10 reads. In addition, bone marrow from rabbit carcasses collected during the same time and location as fly trapping at the Gungahlin site were also positive for RHDV2 recombinants by RT-PCR. This, and the presence of lagoviruses in rabbits and flies in the wider region at that time (Hall et al., 2019), suggests that pathogenic lagoviruses were circulating at the time of sampling and the small number of reads in fly libraries were *bona fide*. In contrast, no legitimate mapping occurred when ectoparasite reads were mapped to a MYXV reference genome (NC_001132.2). This is consistent with the absence of visible clinical signs of myxomatosis in the sampled rabbits. In addition, no viruses with known pathogenic potential in humans were detected in fleas or flies.

## Discussion

A rapidly changing climate increases the potential for ectoparasite-mediated pathogen transmission (Ogden, 2017). A key question is what proportion of the viruses detected in ectoparasites are potentially transmissible to their vertebrate hosts and vice versa, through either the biological or mechanical transmission routes. Similarly, it is important to determine whether some viruses have a greater propensity for mechanical transmission, or a greater capacity to productively infect both vertebrates and invertebrates. The answers to these questions will help reveal the barriers that prevent viruses from evolving vector-borne transmission.

To better understand these key components of vector transmission, we compared the viromes of apparently healthy Australian wild rabbits with those of associated fleas and sympatric flies known to be involved in the transmission of rabbit viruses (Asgari et al., 1998; Sobey & Conolly, 1971). No viruses were found in the lung, liver, duodenum or blood, suggesting the absence of an acute or chronic systemic infection in the wild rabbits sampled for this study. In contrast, considerable viral diversity was detected in the caecal content. This likely reflects the role this organ plays in the digestion of plant matter, such that it is rich in bacteria, other microorganisms, and semi-digested plant material (Forsythe & Parker, 1985; Velasco-Galilea et al., 2018). Based on phylogenetic position, most viruses identified in the caecal content are likely to be associated with the rabbit diet or other commensal microorganisms, such as fungi and protozoa (Figure 2). To our knowledge, equivalent viral meta-transcriptomics analyses on caecal content have not been reported, although an abundance of plant and microorganism-associated viruses is consistent with the faecal viromes of other herbivores (Guan et al., 2018; Woo et al., 2014; Zhang et al., 2017). Importantly, we identified diverse novel viruses in rabbits – *Racaecavirus* and *Orycavirus* - that cluster with other vertebrate-associated viruses (in the *Caliciviridae* and the *Picornaviridae*, respectively) suggesting that the most likely hosts are the rabbits from which they were sampled. In addition, several novel picobirnaviruses were detected, although their true host is uncertain. Overall, the abundance of the potential vertebrate viruses detected in rabbits was relatively low: calicivirus 0.003%, picornavirus 0.025%, picobirnavirus 0.002-0.011%, although benign rabbit viruses have been previously shown to be present at low titre (Capucci, Fusi, Lavazza, Pacciarini, & Rossi, 1996; Strive, Wright, & Robinson, 2009). Furthermore, as these viruses were isolated from caecal content, we would not expect to have sampled a high proportion of rabbit cells and by extension, viruses replicating in these cells.

Members of the *Caliciviridae* and *Picornaviridae* are frequently detected in vertebrates (Shi, Lin, et al., 2018; Zell, 2018), with many cases of confirmed host association (Feinstone, Kapikian, & Purceli, 1973; Ohlinger & Thiel, 1991; Thornhill, Kalica, Wyatt, Kapikian, & Chanock, 1975; Wells & Coyne, 2019). The *Caliciviridae* can be associated with serious illnesses, such as gastroenteritis in humans (Dolin, 1978) and haemorrhagic disease in rabbits (Ohlinger, Haas, Meyers, Weiland, & Thiel, 1990), while the *Picornaviridae* are a diverse group of viruses associated with various diseases in humans and animals (Zell, 2018). Although there are two existing genera that include rabbit caliciviruses, rabbit *Vesivirus* and *Lagovirus,* the novel rabbit calicivirus identified here, *Racaecavirus*, clustered most closely with a pig calicivirus (*St-Valerian swine virus*) and *Marmot norovirus* (Figure 3), both sampled from the gut of healthy animals (L’Homme et al., 2009; Luo et al., 2018). *St-Valerian swine virus* is the only species within the newly classified genus *Valovirus* and the virus identified here (together with *Marmot norovirus*) likely belongs to this genus (L’Homme et al., 2009). The novel rabbit picornavirus we identified in caecal content, *Orycavirus*, was phylogenetically distinct to other rabbit picornaviruses, clustering with enteroviruses and sapeloviruses/sapelo-like viruses (Figure 3). The genus *Enterovirus* includes important human respiratory pathogens, as well as more serious symptoms such as acute flaccid myelitis, meningitis, myocarditis and encephalitis (Wells & Coyne, 2019). Enteroviruses primarily target the gastrointestinal tract and most infections are thought to be asymptomatic (Wells & Coyne, 2019). The genus *Sapelovirus* was initially classified with members from swine, primate and avian hosts, and an unclear link to pathogenicity (Tseng & Tsai, 2007), although the creation of several new genera may now be appropriate (Zell, 2018). The closest relatives of orycavirus were isolated from faeces of apparently healthy cats, bats, and marmots, as well as rodents with unknown disease status (Lau et al., 2011; Lau et al., 2012; Luo et al., 2018). It is notable that the calicivirus and picornavirus detected here cluster with other viruses isolated from the gut content of seemingly healthy vertebrate hosts, tentatively suggestive of a cellular tropism specific to the lower intestinal tract. Additionally, since the sampled rabbits were apparently healthy, the novel calici- and picorna- viruses are likely non-pathogenic. Whether these viruses were present in the founder population of rabbits first introduced into Australia or whether they were exotic incursions awaits additional sampling from diverse locations.

Nine novel species of *Picobirnaviridae* were identified in the rabbit caecum. *Picobirnaviridae* have been detected in several vertebrate species, including rabbits (Ganesh, Masachessi, & Mladenova, 2014; Ludert, Abdul-Latiff, Liprandi, & Liprandi, 1995; Woo et al., 2016), as well as invertebrates (Shi et al., 2016) and diatom colonies (Urayama, Takaki, & Nunoura, 2016). The picobirnaviruses documented here all cluster with the highly diverse and seemingly vertebrate-associated Genogroup 1. The new viruses do not form a monophyletic group by host species (Figure 4), consistent with other members of this family, and diverse picobirnaviruses are commonly found in a single species (Knox, Gedye, & Hayman, 2018; Woo et al., 2016). Consistent with our detection of *Picobirnaviridae* in caecal content, viruses of this family have commonly been isolated from stool samples or cloacal swabs of vertebrates, either with no apparent symptoms or associated with diarrhea (Cummings et al., 2019; Smits et al., 2011; Smits et al., 2012; Woo et al., 2019). Although it has been suggested that these viruses are opportunistic pathogens (Ganesh et al., 2014), the absence of host phylogenetic structure and lack of conclusive detection in solid tissues suggests that vertebrates and invertebrates may not be the true hosts of this virus family. Indeed, based on the presence of conserved prokaryotic ribosomal binding sites, it was recently proposed that prokaryotes are the true hosts of *Picobirnaviridae* (Krishnamurthy & Wang, 2018), which would accord with the lack of taxonomic structure in vertebrate hosts.

A large number of diverse viruses were discovered in fleas collected from rabbits and Calliphoridae, Sarcophagidae and Muscidae flies trapped sympatrically (Figure 1 and Figure 5). Viral composition in ectoparasites varied according to host species (Figure 1) rather than location, consistent with that seen in Australian mosquitos (Shi et al., 2017). The majority of highly abundant viruses were invertebrate viruses, with the remainder likely representing viruses of fungi, protozoa or other commensal microbes (Figure 2). Several viral families/groups identified in rabbit flea libraries were also found in fleas collected from Australian marsupials or rats, including the *Solemoviridae, Iflaviridae*, *Narnaviridae, Phenuiviridae* and *Totiviridae* (Harvey, Rose, Eden, Lawrence, et al., 2019). Generally, viruses from rabbit fleas did not cluster with viruses from other flea species (Figure 2), with the exception of the *Iflaviridae* flea viruses most closely related to Watson virus, a virus of *Pygiopsylla* fleas collected from an Australian marsupial (Harvey, Rose, Eden, Lawrence, et al., 2019). Viruses from six different viral groups/families were identified in both ectoparasites and rabbits (Figure 5), although the ectoparasite viruses were phylogenetically distinct from those found in rabbit caeca (Figure 2). This, and that none of the overlapping viral families were vertebrate-associated, suggests that there may be important barriers to cross-species transmission. Indeed, no highly abundant vertebrate viruses were found in flies or fleas, suggesting that the species investigated here are not likely to be biological vectors for any vertebrate viruses, and that potential arboviruses are rare. Carrion/bush flies and fleas have been implicated in the mechanical transmission of RHDV and MYXV in rabbits (Asgari et al., 1998; Hall et al., 2019; McColl et al., 2002; Sobey & Conolly, 1971). In these cases, no viral replication takes place within the ectoparasite, such that viral abundance would be very low and viral contigs may not be assembled. Accordingly, to detect viruses potentially associated with mechanical transmission, we also explored the low abundant viral reads from the invertebrate libraries (i.e. reads which were not assembled into contigs). This revealed evidence of RHDV and related lagoviruses (*Caliciviridae*) in three Calliphoridae fly species (Figure 6) - a family of flies associated with RHDV transmission (Asgari et al., 1998; Hall et al., 2019; McColl et al., 2002). Since the introduction of RHDV into Australia in 1995, several related viruses have been detected, including recombinants of the original RHDV and RHDV2, or RCV-A1 benign viruses and RHDV2 (Hall et al., 2015; Hall et al., 2018; Mahar, Read, et al., 2018). At least two RHDV2 variants were detected in fly reads (RHDV/RHDV2 and RCV-A1-like/RHDV2), both known to be circulating at that time (Hall et al., 2019; Hall et al., 2018; Mahar, Hall, et al., 2018), and were confirmed by RT-PCR. A small number of RHDV2 reads were also identified in rabbit libraries, and incidentally, RHDV2 was detected by RT-PCR in dead rabbits found synchronously at the study site. Since RHDV infection is generally acute and susceptible rabbits die rapidly (Cooke & Fenner, 2002), RHDV-like reads were likely detected from recovering animals, in which RHDV RNA is detectable for at least 15 weeks post-infection (Gall, Hoffmann, Teifke, Lange, & Schirrmeier, 2007). These results demonstrate that mechanically transmitted viruses can be detected concurrently in the vertebrate host and ectoparasite using a meta-transcriptomics approach, even in the case of highly virulent viruses not known to cause persistent infections. Interestingly, rabbit astrovirus was also detected in *Sarcophaga impatiens* and *Chrysomya varipes*, although no reads were detected in rabbit material. This virus has been associated with enteric disease in rabbits, but may be detected in the gut in the absence of symptoms (Martella et al., 2011). The detection of rabbit astrovirus in flies is of interest as it suggests that astrovirus may be present in Australian wild rabbit populations and must be shed at high titres if it was acquired from faeces. However, as we did not detect any reads in healthy rabbits, more work is clearly needed to establish whether rabbit astroviruses can be transmitted by arthropods.

No viruses known to infect humans, or indeed any other vertebrates besides leporids, were detected in the sampled flies. These fly species are attracted to carrion and faeces, a factor that would promote the mechanical transmission of excreted viruses or those present in carcasses (Norris, 1965). Due to their excessive numbers, rabbit carcasses and faeces are not uncommon in rabbit-infested areas (like the sampling locations), whereas human remains and faeces are hopefully rarer and less accessible. However, we may have expected to find more viruses of livestock (site 1) and native vertebrate species (site 1 and 2), which are abundant in the sampling locations. Hence, vertebrate-associated viral mechanical transmission by fly species may be uncommon, and factors such as high prevalence and high virus load in carcasses or faeces - as seen for RHDV-like viruses - may therefore be necessary for mechanical transmission (Mahar, Hall, et al., 2018; Neimanis, Larsson Pettersson, Huang, Gavier-Widen, & Strive, 2018). In contrast to flies, no vertebrate virus reads - including MYXV - were detected in fleas, although their behaviour of feeding on vertebrate blood rather than carcasses and faeces may limit opportunities for mechanical transmission to periods of acute systemic or viraemic infections. As such, ectoparasite behaviour and host preference, alongside viral pathogenesis and prevalence, are likely important for mechanical transmission.

In sum, while rabbits and ectoparasites carry viruses from some of the same viral families, viruses from ectoparasites are phylogenetically distinct from viruses found in rabbit caecal content, suggesting that major host barriers exist that prevent invertebrate viruses from establishing productive replication cycles in vertebrates. Importantly, however, flies carried a very low abundance of vertebrate viruses with pathogenic capacity in rabbits, including RHDV for which fly-mediated mechanical transmission has already been demonstrated. Hence, although biological transmission appears difficult to evolve, flies may serve as important mechanical vectors for rabbit-associated viruses.

## Acknowledgements

We thank Tegan King and Oliver Orgill and team from the ACT Parks and Conservation Services for sample acquisition, and Nina Huang for detection of lagoviruses in rabbit bone marrow. ECH is supported by an ARC Australian Laureate Fellowship (FL170100022). The authors acknowledge the Sydney Informatics Hub and the University of Sydney’s high-performance computing cluster Artemis for providing the high-performance computing resources that have contributed to the research results reported within this paper.

## Data accessibility

- Raw data: NCBI SRA BioProject XXXX.
- Viral contigs presented in phylogenies: GenBank accession numbers XXXXX-XXXX.

## Author contributions

JE Mahar was involved in sample collection and research design, performed research, analysed data, and wrote the paper. M Shi contributed analytical pipelines and was involved in research design and manuscript editing. RN Hall was involved in sample collection, fly identification, research design and manuscript editing. T Strive was involved in research design, sample collection and manuscript editing. EC Holmes designed and obtained funding for research, assisted with data analysis and manuscript writing.

