## Supplementary Table 1 for "Limited overlap in RNA virome composition among rabbits and their ectoparasites reveals barriers to virus transmission"

**Table S1. Details of the virus sequence alignments used in the phylogenetic analyses**

| Virus family/group | RNA virus type | Alignment length† | No. sequences in alignment |
|---|---|---|---|
| *Caliciviridae* | Positive-sense | 397 | 40 |
| *Picornaviridae* | Positive-sense | 412 | 114 |
| *Picobirnaviridae* | Positive-sense | 468 | 99 |
| *Solemoviridae* | Positive-sense | 303 | 109 |
| *Tombusviridae* | Positive-sense | 356 | 113 |
| *Iflaviridae* | Positive-sense | 448 | 47 |
| *Narnaviridae* | Positive-sense | 304 | 107 |
| *Dicistroviridae* | Positive-sense | 419 | 32 |
| *Flaviviridae* | Positive-sense | 301 | 69 |
| *Virgaviridae/Bromoviridae* | Positive-sense | 385 | 75 |
| *Nodaviridae* | Positive-sense | 403 | 63 |
| *Hypoviridae* | Positive-sense | 258 | 17 |
| *Orthomyxoviridae* | Positive-sense | 362 | 46 |
| *Monovirales/Chuviridae* | Positive-sense | 526 | 135 |
| *Bunyavirales* | Negative or ambisense | 475 | 145 |
| *Totiviridae/Chrysoviridae* | Double-stranded | 387 | 103 |
| *Reoviridae* | Double-stranded | 444 | 97 |
| *Partitiviridae* | Double-stranded | 270 | 110 |

†Alignment length refers to the final length of the alignment in amino acids, after trimming with Trimal
